# Association between the brain-derived neurotrophic factor Val66Met polymorphism and overweight/obesity in Mexican pediatric population

**DOI:** 10.1101/215798

**Authors:** José Darío Martínez-Ezquerro, Mario Enrique Rendón-Macías, Gerardo Zamora-Mendoza, Jacobo Serrano-Meneses, Beatriz Rosales-Rodríguez, Deyanira Escalante-Bautista, Maricela Rodríguez-Cruz, Raúl Sánchez González, Yessica Arellano-Pineda, Mardia López-Alarcón, María Cecilia Zampedri, Haydeé Rosas-Vargas

**Affiliations:** Unidad de Investigación Médica en Genética Humana, Hospital de Pediatría, Centro Médico Nacional Siglo XXI, Instituto Mexicano del Seguro Social (IMSS), Ciudad de México, Distrito Federal, México; Posgrado en Ciencias Biológicas, Universidad Nacional Autónoma de México (UNAM); Av. Ciudad Universitaria 3000, C.P. 04510, Coyoacán, Distrito Federal, México; Unidad de Investigación en Epidemiología Clínica, Hospital de Pediatría, Centro Médico Nacional Siglo XXI, Instituto Mexicano del Seguro Social (IMSS), Ciudad de México, Distrito Federal, México; Unidad de Investigación Médica en Nutrición, Hospital de Pediatría, Centro Médico Nacional Siglo XXI, Instituto Mexicano del Seguro Social (IMSS), Ciudad de México, Distrito Federal, México; Laboratorio de Genómica Funcional, Instituto Nacional de Medicina Genómica (INMEGEN), Ciudad de México, Distrito Federal, México

**Keywords:** BDNF, BMI z-score, body weight, overweight, obesity

## Abstract

**Background:** The functional brain-derived neurotrophic factor (BDNF) rs6265 (G196A; Val66Met) single nucleotide polymorphism has been associated with eating disorders, BMI and obesity in distinct populations, both adult and pediatric, with contradictory results involving either Val or Met as the risk variant.

**Aim of the study:** To determine the association between the BDNF Val66Met polymorphism and BMI in Mexican children and adolescents.

**Methods:** BDNF Val66Met genotyping by restriction fragment length polymorphism and nutritional status characterized by their BMI-for-age z-scores (BAZ) from pediatric volunteers recruited in Mexico City (n=498) were analyzed by Fisher’s exact test association analysis. Standardized residuals (R) were used to determine which genotype/allele had the major influence on the significant Fisher’s exact test statistic. Odds ratios were analyzed to measure the magnitude and direction of the association between genotype and normal weight (≥ −2 SD < +1 SD) and overweight (≥ +1 SD, including obesity, Ow+Ob) status with 95% confidence intervals to estimate the precision of the effect as well as 95% credible intervals to obtain the most probable estimate.

**Results:** Comparisons between GG (Val/Val), GA (Val/Met) and AA (Met/Met) genotypes or Met homozygotes vs. Val carriers (combination of GG and GA genotypes) showed significant differences (p=0.034 and p=0.037, respectively) between normal weight and the combined overweight and obese pediatric subjects. Our data showed that children/adolescents homozygous for the A allele have increased risk of overweight compared to the Val carriers (Bayes OR= 4.2, 95% CI**[1.09-33.1]).

**Conclusion:** This is the first study showing the significant association between the BDNF rs6265 AA (Met/Met) genotype and overweight/obesity in Mexican pediatric population.

- Mexico has one of the highest pediatric overweight and obesity prevalence
- BDNF has been associated with body weight regulation.
- The BDNF rs6265 SNP (G196A; Val66Met) has been associated with eating disorders, BMI and obesity, with contradictory results in both adults and children.
- We found significant associations between BDNF Val66Met AA (Met/) genotype and overweight/obesity in Mexican pediatric population
- Met homozygote children/adolescents increased four times the risk of being classified in the overweight group (Ow+Ob) relative to Val carriers

## Introduction

### Obesity epidemic

Obesity is a global increasing epidemic for both children (1) and adults (2) that compromises human well being. The most critical comorbidities related to adipose tissue excess range from rheumatological conditions to type 2 diabetes mellitus, cardiovascular disease, and increased risk of cancer (3). According to the World Health Organization (WHO) (4), overweight and obesity currently affects 1.9 billion adults and 41 million children under the age of five all around the world, accounting as the fifth leading risk for global deaths with at least 2.8 million adults dying each year as a result of these conditions. Up to the past decade, developing countries such as Mexico, China and Thailand have had the most dramatic increase in obesity (5). Recently, it was published that Mexico has the second prevalence of obesity in the adult population with 22 million obese (30%) in addition to the 26 million adults with overweight, while ranking fourth in children (6), with an overall overweight (Ow) or obesity (Ob) prevalence of 28.8% in children <19 years of age as for the most recent results from the Mexican health and nutritional survey (ENSANUT, 2012) (7,8). In the last 24 years, the highest prevalence was observed among children and adolescents living in urban areas and those from the highest socioeconomic level, while the rate of increase was higher in the lowest socioeconomic status (8).

In general, it is assumed that obesity results from a combination of genetic susceptibility, increased availability and consumption of high-energy foods as well as a decreased requirement and performance of physical activity as a consequence of modern life styles (9). Obesity is a complex condition determined by an intricate interplay of genetic and environmental factors (10). Genetic variants are estimated to account for a range between 40 to 70% of the heritability of BMI (11,12), including single mutations as well as single nucleotide polymorphisms (SNPs) causing from severe impairment in appetite regulation and early-onset overweight to slightly increased BMI or early-onset obesity (11).

### BDNF and obesity

As for genetic susceptibility, known single-gene mutations (13,14) or syndromes (15) may explain only a small fraction (∼5%) of childhood-onset obesity. However, as mentioned previously, obesity can mainly be the result of the imbalance between caloric intake and energy expenditure, so by studying the genes involved in appetite regulation we will be able to unravel the essential molecular network involved in obesity.

One such molecule that has been associated with body weight regulation is the brain-derived neurotrophic factor (BDNF). BDNF is a member of the neurotrophin family of small secreted proteins with major roles in central nervous system (CNS) development. Current data from Ensembl shows that BDNF is located at locus 11p14.1, extends over approximately 67 kb, contains 12 exons with 9 functional promoters for tissue and brain-region specificity and originates 19 transcripts by alternative splicing (16). Information about the pro-BDNF proteolytic processing, mature BDNF and its receptors p75^NTR^ and Trkb, respectively, be reviewed elsewhere (17).

Although it is widely expressed among several tissues (18), BDNF is abundant in the CNS (19,20), predominantly in the hippocampus, amygdala, cerebral cortex, and hypothalamus (21-23). BDNF plays a critical role in nervous system development and function (24,25), and particularly, exerts an anorexigenic function in the brain (26). BDNF molecular alterations have been implicated in conditions affecting body weight such as eating disorders (27,28). One of these variations affecting BDNF is the Val66Met single nucleotide polymorphism (G196A; SNP rs6265). In particular, the 66Met (A variant) allele is biologically relevant as it alters the intracellular processing, trafficking and activity-dependent secretion of BDNF (29,30), and has been associated with several clinical traits such as early seizures, bipolar affective disorders, obsessive-compulsive disorders, eating disorders, BMI, and obesity (17).

As with adults (31), studies involving children and adolescents attempting to examine the association between the BDNF rs6265 polymorphism and age-and-sex specific nutritional status characterized by their BMI-for-age z-scores (BAZ) have shown contradictory results. Some of them have found association between this SNP and childhood BAZ at the upper tail of the BMI distribution in children with European ancestry (32), as well as for BMI and obesity in Chinese (33-35), European American (36), and Croatian (37) children; while others reported no association with BMI in Spanish (38), with BAZ in Mexican children (39), and with extreme obesity in German children and adolescents (40).

Although the BDNF 66Met (A allele) presents greater plausibility of being associated with BMI increase and overweight/obesity as it is a functional variant that generates subcellular translocation and activity-dependent secretion deficiencies of BDNF which could resemble the BDNF deficiencies associated with obesity (41,42), several articles have pointed to the Val66 allele (G variant) as the risk allele associated with BMI or obesity risk (33-35,43), while others point to the Met66 allele (A variant) (37,44,45), and even to the heterozygous genotype AG (37,46). In example, it has been observed in German children that 66Met carriers, although associated with lower BMI, had an increased calorie intake and reported higher carbohydrates and proteins consumption (47), while in Chinese children carriers of the A allele are at increased risk of obesity when moderate to low physical activity levels are reported (45).

At present, association studies involving BDNF rs6265 and BMI in children and adolescents are still scarce and conflicting. Therefore, the aim of this study was to analyze the relationship and determine the association between the BDNF Val66Met polymorphism and nutritional status characterized by their BMI-for-age z-scores (BAZ) in Mexican pediatric subjects.

## Material and methods

### Subject recruitment and sample collection

Samples were obtained from Mexican pediatric volunteers (n=498), 282 girls (56.6%) and 216 boys (43.4%) between 5-17 years old (312 overweight children and 186 normal weight controls) without any metabolic condition reported. This study was performed at Unidad de Investigación Médica en Genética Humana (UIMN), Hospital de Pediatría, at Centro Médico Nacional Siglo XXI from Instituto Mexicano del Seguro Social (CMN Siglo XXI, IMSS). Our protocol was reviewed and approved by the Ethical Committee of IMSS and assigned with the registry number R-2009-3603-9; both children and parents provided written informed consent for participation in the study before any study-related procedures were performed.

### Biological parameters

Nutritional status categories for children and teenagers were diagnosed by calculating both their individual BMI as weight(kg)/height^2^(m^2^) as well as their BMI-for-age z-scores for either girls or boys (BAZ), following WHO’s growth reference data for 5-19 years (48) and employing the WHO AnthroPlus software (WHO 2007 R macro package) (49). Participants were then classified in two main BAZ categories: normal weight (> −2 SD < +1 SD; n= 186) and overweight group, including overweight and obesity (≥ +1 SD; n= 312). Only BAZ values between −3 and +5 z-scores were considered valid and included in this analysis (7).

### Genotyping

Five milliliters of blood from each fasting participant were collected by a standard method in an EDTA tube. For the DNA preparation a commercial kit (Illustra blood genomicPrep Mini Spin Kit, GE Healthcare) was used.

The BDNF Val66Met SNP rs6265 genotype (G196A) was obtained, as previously described (50), using a polymerase chain reaction-restriction fragment length polymorphism (PCR-RFLP) method with forward (5’-ACTCTGGAGAGCGTGAAT-3’) and reverse (5’-ATACTGTCACAC ACGCTC-3’) primers, and further digestion of the PCR product with NlaIII enzyme (Cat. No. R0125S, New England Biolabs). From the five possible restriction fragments for this Val66Met amplicon, the genotype was identified by the size and distribution of three bands: 243-bp for the G variant (Val), 168-bp and 75-bp bands for the A variant (Met), and these three bands for GA heterozygotes (Val/Met), on 2.5% (w/v) agarose gel electrophoresis. A random selection of 15 samples was performed for validation with genomic DNA sequencing.

### Statistical analysis

The data analysis was carried out with free and commercial software, R and SPSS, respectively as well as online tools for statistical computation and visualization (51,52). Baseline characteristics for quantitative variables are presented as arithmetic mean and standard deviation (mean ± SD), and evaluated using one-way analysis of variance (ANOVA), while simple frequencies (n) and percentages (%) were used for qualitative variables.

Allele and genotypic frequencies were analyzed for compliance with chi-square Hardy-Weinberg equilibrium (HWE) with an online calculator for biallelic markers that includes an analysis for ascertainment bias for dominant/recessive models due to biological or technical causes (52) for both the whole pediatric sample as well as for each of the two BAZ categories.

To evaluate the possible association between the BDNF rs6265 polymorphism and BAZ nutritional status, genotype frequencies of the pediatric population were compared against the two main nutritional categories: normal weight and overweight, including obesity. To determine if there were significant differences in the frequency of occurrence for each genotype: GG (Val/Val), GA (Val/Met), and AA (Met/Met) or allele (Val or Met) in a particular nutritional group, we performed Chi-square (χ^2^) statistics and Fisher’s exact test. Standardized residuals (R) were obtained as a measure of the strength of the difference between observed and expected values to determine how significant the frequencies are to the chi-square (χ^2^) value and which frequencies had the major influence on the significant chi-square (χ^2^) test statistic; positive or negative standardized residuals indicate that there are more or less than expected, respectively. Alternatively, the AA (Met/Met) and GA (Val/Met) groups were combined into Met carriers and compared against the homozygous GG group (Val homozygotes). Both the HWE and the frequency of each genotype according to their classification into a BAZ group were plotted in *de Finetti diagrams* to visualize the proportions and possible significant deviations from HWE of the bi-allelic marker as implemented in the program DeFinetti (53) as well as to explore the clustering trends between genotypes and BAZ categories (54).

To establish the magnitude of the association between Val66Met genotypes and nutritional groups, we performed logistic regression analysis considering genotypes or alleles as independent variables and BAZ categories as dependent variables comparing Normal weight vs. Overweight and Obesity combined (Ow+Ob). Odds ratios (ORs) with 95% confidence intervals (CI 95%) were obtained for precision. All results were considered statistically significant when two-tailed Fisher’s exact test p-value was < 0.05.

A bayesian analysis was performed with JASP using default priors (55) to determine the relative plausibility of the data under the null hypothesis versus the alternative through the Bayes factor (BF), when comparing the null hypothesis (H0) of no association between Val66Met genotypes and BAZ, as well as the alternative hypothesis (H1) as the association between them. In addition, the bayesian OR with a 95% credible interval (CI**) was obtained.

### Results

We recruited 498 Mexican children and adolescents attending the Unidad de Investigación Médica en Nutrición from the Hospital de Pediatría at CMN Siglo XXI, IMSS (Mexico City). The phenotypic characteristics and BDNF Val66Met genotypes of the pediatric participants, 282 girls (56.6%) and 216 boys (43.4%), ranging between 5 and 17 years (mean age: 12.2 ± 2.02 years) are shown in Table I. According to the BAZ categories, 37.3% of the participants had normal weight and 62.6% were overweight (corresponding to 26.1% overweight and 36.5% obesity). The BDNF Val66Met allele frequencies were 0.85 and 0.15 for G (Val) and A (Met), respectively, while the genotype frequencies were 71.7% for GG (Val/Val, n=357), 25.9% for GA (Val/Met, n=129), and 2.4% for AA (Met/Met, n=12). Hardy-Weinberg equilibrium (HWE) criteria under a model of ascertainment was met (χ^2^ =0.007; df=1; p=0.93), indicating no deviation from the Hardy-Weinberg equilibrium in this study, and that our data had no gain/losses bias in the genotype counts. There were no differences in genotype distribution between female and male subjects (Table I). Our results are similar to those reported previously for other pediatric populations (56-58), and as expected under HWE.

**Table I.** Baseline phenotypes and BDNF Val66Met (G196A) genotypes of Mexican children and adolescents (n=498) ranging between 5-17 years.

| Phenotypes | Participants |  | Frequency n (%) |  |
| --- | --- | --- | --- | --- |
| Age (mean ± SD years) | 12.2 ± 2.02 |  | 498 (100) |  |
| Sex | Female (girls) |  | 282 (56.6) |  |
|  | Male (boys) |  | 216 (43.4) |  |
| Nutritional Status (BAZ) | Normal weight |  | 186 (37.3) |  |
|  | Overweight |  | 130 (26.1) |  |
|  | Obesity |  | 182 (36.5) |  |
|  |  |  | (%) within gender** |  |
|  | Genotype | Population* | Female | Male |
| BDNF Val66Met | GG (Val/Val) | 357 (71.7) | 209 (74.1) | 148 (68.5) |
|  | GA (Val/Met) | 129 (25.9) | 67 (23.8) | 62 (28.7) |
|  | AA (Met/Met) | 12 (2.4) | 6 (2.1) | 6 (2.8) |

\* HWE: $\chi^2 = 0.007$ , df=1 p=0.93; \*\* $\chi^2 = 1.9$ , df=2 p=0.38
BAZ, BMI-for-age z-scores

Figure 1 illustrates the *de Finetti* distributions (53,54) of the BDNF rs6265 polymorphism according to (A) the test for deviation from HWE of both the normal weight and overweight group (HWE: p=0.053 and p=0.124 with Fisher’s exact test, respectively), as well as (B) the frequency and location of the pediatric sample according to both their genotypes and BAZ classification. These visualizations showed that not only the whole sample but each of the two main BAZ groups, normal weight and overweight group, are in HWE, but also that there is a clear trend towards clustering closer to the overweight or obesity groups when bearing the AA genotype. GA (n=129) and AA (n=12) are shown with green and orange circles, and red dot, respectively, in proportion with the observed counts per BAZ category.

**Figure 1.**
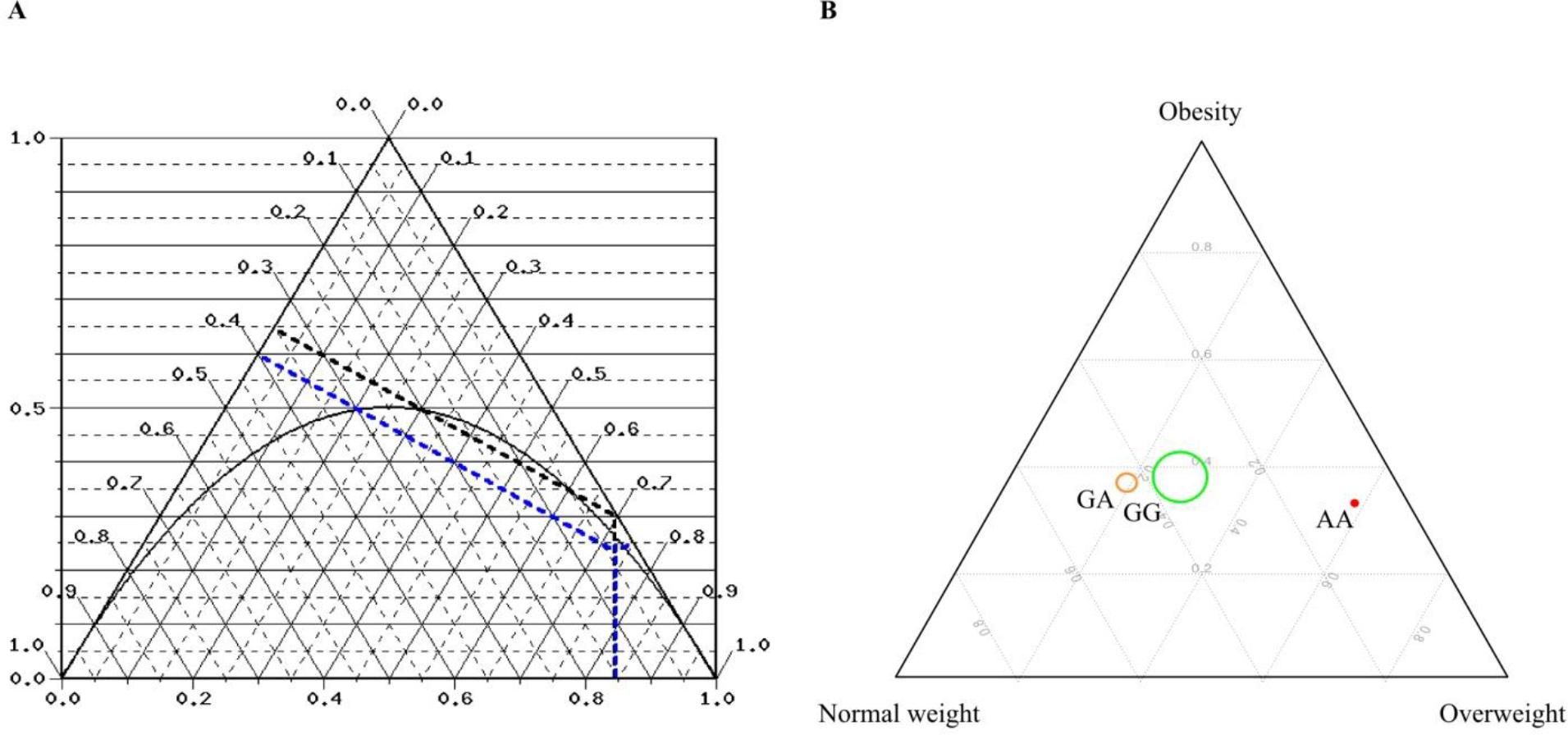
De Finetti ternary diagrams for Mexican pediatric population (n=498) of BDNF rs6265. These visualizations represents both the Hardy-Weinberg equilibrium of the bi-allelic marker in the pediatric population as well as their clustering into normal weight (> −2 SD < +1 SD), overweight (≥ +1 SD < +2 SD) and obesity (≥ +2 SD) phenotypes according to their BDNF rs6265 genotype. (A) Ternary diagram for the single nucleotide polymorphism with a Hardy-Weinberg parabola for normal weight (black dotted line, p=0.053) and overweight groups (≥ +1 SD; blue dotted line, p=0.124) (https://ihg.gsf.de/cgibin/hw/finetti2.pl). (B) Ternary plot visualizing the location and frequency of genotypes in relation to nutritional status classified by WHO’s BAZ cut-offs. The genotypes GG (n=357), GA (n=129) and AA (n=12) are shown with green and orange circles, and red dot, respectively, in proportion with the observed counts per BAZ category.

The aim of this study was to determine a possible association between BDNF rs6265 and nutritional status. We observed significant differences between the frequencies of the BDNF GG (Val/Val), GA (Val/Met) and AA (Met/Met) genotypes (p=0.034), as well as between the Met homozygotes in comparison to the combined Val/Val and Val/Met genotypes grouped into the Val carriers (p=0.037) depending on the normal weight or overweight status (Table II). The frequency of the Val and Met alleles or the Met carriers (the combined Val/Met and Met/Met genotypes in comparison to the homozygous Val/Val genotype) did not show statistical differences. Although the absolute standardized residuals (R) values were not ≥ 1.96 which would indicate that the frequency of that group is significantly contributing to the difference between proportions, both the lowest and highest R values were detected for the AA genotype (R= −1.6 and +1.3) in the normal weight and overweight group (Ow+Ob), respectively, when compared against GG and GA. Similar results were obtained for the lowest and highest R values from Met homozygotes (R= −1.4 and 1.1) when compared against the Val carriers in the normal weight and Ow+Ob group, respectively. These results indicate that the significant difference between proportions is mainly the result of the lower and higher frequency of the AA (Met/Met) genotype among subjects in the normal weight (0.5%) and overweight group (3.5%), respectively (Table II).

**Table II.** The BDNF genotype and allele count and frequencies (%) in Mexican children and adolescents (n = 498) subdivided into two main BAZ groups, normal weight (< -2 > +1 SD) and overweight group (> +1 SD, including overweight and obesity)

| BDNF Val66Met genotype | Normal weight<br>n (%) | Overweight group<sup>a</sup><br>n (%) |
| --- | --- | --- |
| GG (Val/Val) | 129 (69.4) | 228 (73.1) |
| GA (Val/Met) | 56 (30.1) | 73 (23.4) |
| AA (Met/Met) | 1 (0.5) | 11 (3.5) |
| Fisher's exact test **p=0.034** |  |  |
| R= -1.6 and +1.3 for AA genotype in the normal and overweight groups, respectively |  |  |
| Val allele | 314 (84.4) | 529 (84.8) |
| Met allele | 58 (15.6) | 95 (15.2) |
| Fisher's exact test p=0.93 |  |  |
| Val carriers | 185 (99.5) | 301 (96.5) |
| Met homozygotes | 1 (0.5) | 11 (3.5) |
| Fisher's exact test **p=0.037** |  |  |
| R= -1.4 and +1.1 for Met homozygotes in the normal and overweight groups, respectively |  |  |
| Met carriers | 57 (30.6) | 84 (26.9) |
| Val homozygotes | 129 (69.4) | 228 (73.1) |
| Fisher's exact test p=0.411 |  |  |

<sup>a</sup> $\geq +1$ SD, including overweight and obese children; R=absolute value of the standardized residual

Given the contradictory results reported for the BDNF rs6265 SNP in pediatric as well as adult populations in which either variant has been associated with BMI or Ow+Ob-related conditions, we explored several genotype combinations considering either the A or G variant as the risk allele to obtain the directions of the effects. Our results showed that when considering the A variant as the risk allele, the AA (Met/Met) genotype increased more than six times the risk of being classified within the overweight group (Ow+Ob) when compared against each genotype: GG (6.22, p=0.039), AG (8.43, p=0.015), or both GA+GG (6.76, p=0.028). On the other hand, when considering the G variant as the risk allele, several combinations (GG vs AA, GA vs AA and GG+GA vs AA) showed that the common variant G had a significant protective effect against being classified in the overweight (Ow+Ob) group (Figure 2A).

**Figure 2.**
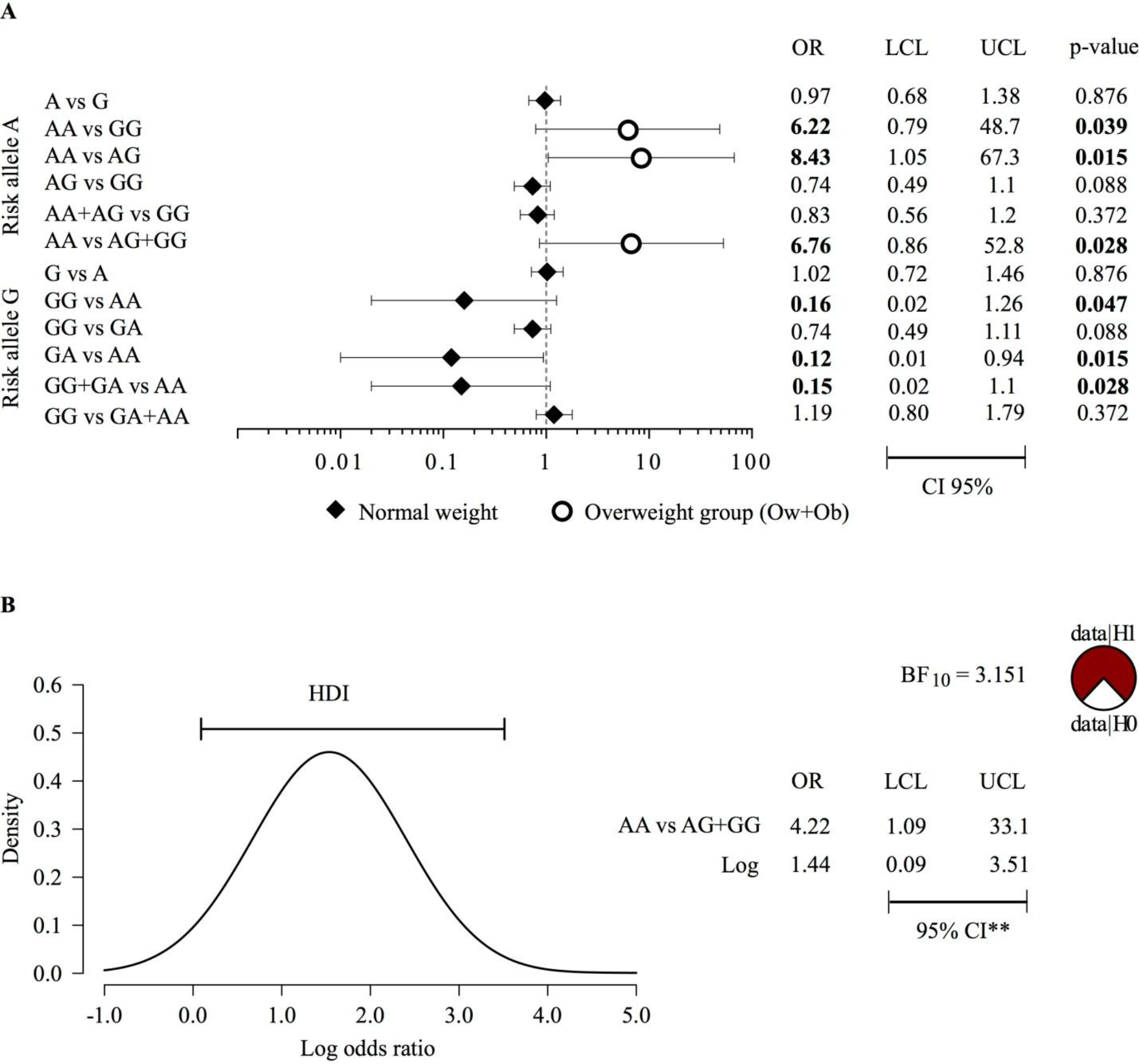
Risk estimation between BDNF rs6265 genotypes and BAZ categories in Mexican pediatric population. The image shows (A) a forest plot visualizing the magnitude and precision of the association of the effect of either allele A or G at rs6265 in BDNF on overweight and obesity combined, and (B) the 95% highest density interval for the association between AA vs GG+GA when compared against the BAZ groups, where every point inside the area under the curve between the limits has higher credibility (probability density) than any point outside the interval. The pie chart visualize the Bayes factor where the red predominance over white indicate evidence for the alternative hypothesis. BAZ, BMI for-age-and-sex z-scores; OR, odds ratio; CI 95%, 95% confidence interval; LCL, lower confidence interval; UCL, upper confidence interval; Ow, overweight; Ob; obesity. ORs and p-values in bold indicate statistical significance by two-tailed Fisher’s exact test. BF_10_, Bayes factor for the alternative hypothesis; H0, null hypothesis; H1, alternative hypothesis; HDI, highest density interval; 95% CI**, 95% credible interval. Both X-axis represents values at Log10 scale.

Further, to estimate a more credible OR we performed a Bayesian analysis. Results of the bayesian analysis are only shown for the association between AA vs GG+GA when compared against the BAZ groups (Figure 2B). First, to quantify the evidence provided by the observed data in favor of one hypothesis over the other we calculated the Bayes factor (BF), where values >1 indicate evidence in favor of the alternative hypothesis (H1: the probability of AA (Met/Met) genotype is higher in the overweight group). The Bayes factor for the alternative hypothesis (BF_10_) was 3.151 suggesting that these data are 3.151 more likely to be observed under the alternative hypothesis (H1). The bayesian OR for this association was 4.22, indicating the most probable associated risk value for Ow+Ob with a 95% credible interval of 1.09 - 33.1.

## Discussion

To our knowledge, this is the first study in Mexican pediatric population showing a significant association between the BDNF Val66Met (rs6265) Met homozygous and nutritional status characterized by their BMI-for-age z-scores (BAZ).

According to an update of our previous bibliometric analysis for BDNF Val66Met in three main databases: Web of Science, Pubmed and Scopus (59), a previous study in healthy Mexican school-aged children between 6-15 years of age recruited from a summer camp (60) did not find association between this SNP and BMI for-age z-scores (39); however, the authors analyzed this association as a linear function of BAZ, instead of considering nutritional status as well established overweight and obesity categories. In addition, they considered the G variant as the risk allele and only reported the risk allele frequency, but not the frequency of the AA (Met/Met) genotype, which could have allowed a comparison between our samples. At least across 58 global populations, the derived Met allele exhibits a wide allelic distribution variability ranging from 0 to 72% (61); in our sample, this frequency is 15.2%.

Given the low frequency of the AA (Met/Met) genotype in our sample, employing BAZ categories resulted in the increase of statistical power, allowing us to detect its association with overweight and obesity. Indeed, WHO’s standards were chosen since they depict normal (non-obese) childhood growth and can be used to assess children, regardless of ethnicity, socioeconomic status and type of feeding. In addition, it has been reported as a more sensitive criterion to identify overweight and obesity than CDC and IOTF recommendations since they derived from more recent data in which the BMI distribution of the reference populations has shifted towards the right due to an increase in BMI (62-64). Not only the frequency of the AA (Met/Met) genotype in our population is similar to that observed in Croatian children (37), 2.4% and 3%, respectively, but also its association with overweight/obesity. Although difficult to explain, discrepancies observed in the literature about the risk allele associated with BMI, overweight, and obesity may be the result of differences in the clinical criteria for patients selection, allele distribution between populations, strategies for data processing and analysis (ie. BMI-for-age or nutritional status, national or international BAZ cut-offs), gene-environment factors, and the effect of other genes. For example, it has been shown that the effect of several SNPs associated with BMI, overweight and obesity in Europeans does not replicate completely in Mexican children, implying that distinct population genetic susceptibility variations may account for these outcomes (65), in the context of particular gene-environment or gene-diet interactions which would require further exploration.

From the beginning, we hypothesized that the association between the AA genotype with the overweight BAZ category was a more plausible outcome given the clear functional effects of this SNP, in which in an homozygous state the molecular function would be completely altered, while compensated in an heterozygous state. As mentioned previously, the 66Met variant reduces the expression of BDNF, which inhibits excessive calorie uptake and promotes energy expenditure; then, the 66Met variant impairs the normal function of these systems with direct impact in either BMI or Ow+Ob-related outcomes (17). We tested this hypothesis by considering either the A or G allele as the risk variant. As biologically expected, we observed that the significant risk of being overweight (Ow+Ob) was observed only in participants bearing the AA (Met/Met) genotype (Figure 3). It is worth mention that the only children with the AA genotype classified in the normal weight group had a BMI-for age z-score= 0.96, which is almost in the cut-off point for the overweight group (≥ +1 SD) classification. In contrast, either one or two copies of the Val allele, showed a significant protective effect against overweight (Figure 2A).

**Figure 3.**
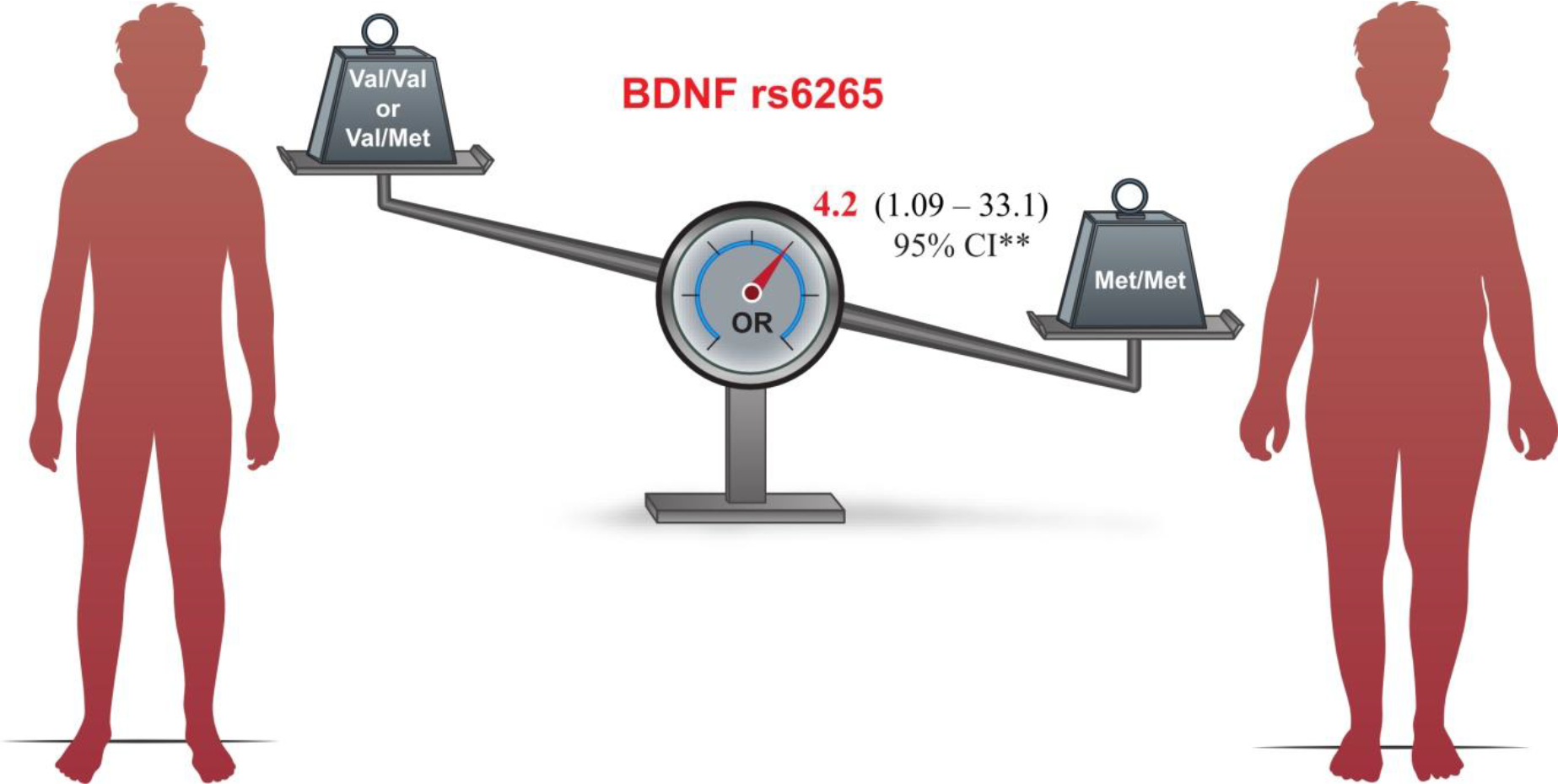
BDNF Val66Met is a genetic risk factor for overweight and obesity in Mexican children and adolescents. The AA (Met/Met) genotype relative to Val carriers (GG and GA genotypes) increased four times (Bayes OR= 4.22, 95% CI**[1.09 – 33.1]) the risk of overweight, including obesity. OR, odds ration; CI**, credible interval.

Our results are partially in line with recent studies that have shown a significant association between obesity (BMI percentile) for Caucasian children and adolescents of the same ethnic (Croatian) background and one or two Met alleles of the BDNF Val66Met polymorphism, each of them increasing their BMI, and with a significant risk for obesity in children bearing the Val/Met genotype (37). However, no significant differences in the distribution of the BDNF Met carriers compared to Val homozygotes were observed for adults from the same ethnic (Croatian) background for normal weight, overweight and obese categories, neither gain or changes during a 35 years of follow-up with three time check-up periods (43.4 ± 4.4, 53.4 ± 4.5, and 77.2 ± 4.5 years; mean age in years ± SD for each period) (66). A recent systematic review and meta-analysis assessing the association of BDNF polymorphisms and BMI, as a representative index of overweight and obesity, has concluded that the rs6265 SNP can be considered as a genetic determinant of obesity (31). Nevertheless, a previous analysis from data of the Brain Resource International Database showed trends towards a lower BMI in adults from 18 to 82 years bearing the Met/Met genotype compared to the Val/Val and Val/Met genotypes as well as when comparing the Met homozygotes to the Val carriers (67).

According to these observations, it results intriguing the possibility that subjects with Met/Met genotype may shift their BMI according to their life span, in which young Met/Met subjects have an increased probability of showing a higher BAZ which will be attenuated in adulthood until shifting towards a lower BMI in old age.

Factors and mechanisms involved in this Met homozygous-BMI shift still remain to be elucidated. In fact, follow-up studies considering changing eating patterns, physical activity or sedentary behaviors, differential mechanisms and distinct combination of factors across lifespan regulating food intake and caloric expenditure between children, adults, and elderly subjects from distinct populations must be considered. Clearly, brain-regulating hormonal signals like leptin and insulin (68), with physiological effects particularly in the hippocampus, amygdala, cerebral cortex, and hypothalamus, brain regions with abundant BDNF expression (21-23), and some of them involved in weight regulation and food intake via its actions on specific hypothalamic nuclei (69) should be considered in further molecular combinatorial analysis, as the homeostatic imbalance of these nutritional signals has been associated to weight loss in lean older adults (70).

## Conclusion

Finally, we have confirmed BDNF Val66Met, particularly the AA genotype (Met variant), as a genetic risk factor for nutritional BAZ status in Mexican children and adolescents. However, studies on this polymorphism in Mexican pediatric population should be replicated in a larger sample and include the variables mentioned above that may be involved in the association discrepancies reported among populations. Further research should be encouraged towards BDNF and functional variants such as Val66Met associated with energy metabolism, food regulation and BMI, particularly in countries like Mexico widely affected by this health-threatening condition.

## Acknowledgements

JDME thanks the Posgrado en Ciencias Biológicas, Universidad Nacional Autónoma de México (UNAM) for the formation received during his PhD studies in Biological Sciences. This article is a requirement to obtain the PhD degree by Posgrado en Ciencias Biológicas, UNAM.

This research was funded by Fondo de Investigación en Salud (FIS) from the Instituto Mexicano del Seguro Social (IMSS) FIS/IMS/PROT/1087. JDME was supported with a scholarship from Consejo Nacional de Ciencia y Tecnología (Conacyt; CVU/Becario: 412887/263531).

The authors thank Mauricio Villagrán-Rendón for his help with the figure design.

## Author contributions

JDME, GRZM, JSM, YAP, MRC, RSG, MLA and MCZ: collection and/or assembly of the data; JDME, GRZM, BRR, and DEB: sample processing and genetic analysis; JDME and MERM: statistical analysis and data interpretation; JDME: data analysis and visualization, and manuscript writing; HRV: conception, design and financial support of the study, revised the article and approved the final version

## Abbreviations

BDNF: brain-derived neurotrophic factor
WHO: World Health Organization
CNS: central nervous system
RFLP: restriction fragment length polymorphism
SNP: single nucleotide polymorphism
BAZ: BMI-for-age z-score
SD: standard deviation
Ow: overweight
Ob: obesity
OR: odds ratio
CI: confidence interval
CI**: credible interval

